# Inverse Finite Element Framework Combining Ultrasound Imaging and Inflation Testing of PVA Artery Phantoms

**DOI:** 10.1101/2024.05.01.591942

**Authors:** Yasmine Guendouz, Noor Adeebah Mohamed Razif, Floriane Bernasconi, Gordon O’ Brien, Robert D. Johnston, Caitríona Lally

## Abstract

The clinical decision to establish if a patient with carotid disease should undergo surgical intervention is primarily based on the percent stenosis. Whilst this applies for high-grade stenosed vessels (>70%), it falls short for other cases. Due to the heterogeneity of plaque tissue, probing the mechanics of the tissue would likely provide further insights into why some plaques are more prone to rupture. Mechanical characterisation of such tissue is nontrivial, however, due to the difficulties in collecting fresh, intact plaque tissue and using physiologically relevant mechanical testing of such material. The use of polyvinyl alcohol (PVA) is thus highly convenient because of its acoustic properties and tuneable mechanical properties.

**Methods:** The aim of this study is to demonstrate the potential of polyvinyl alcohol phantoms to simulate atherosclerotic features. In addition, a testing and simulation framework is developed for full PVA vessel material characterization using ring tensile testing and inflation testing combined with non-invasive ultrasound imaging and computational modelling.

**Results:** Strain stiffening behaviour was observed in PVA through ring tensile tests, particularly at high freeze-thaw cycles. Inflation testing of bi-layered phantoms featuring lipid pool inclusions demonstrated high strains at shoulder regions. The application of an inverse finite element framework successfully recovered boundaries and determined the shear moduli for the PVA wall to lie within the range 27 kPa to 53 kPa.

**Conclusion:** The imaging-modelling framework presented facilitates the use and characterisation of arterial mimicking phantoms to further explore plaque rupture. It also shows translational potential for non-invasive mechanical characterisation of atherosclerotic plaques to improve the identification of clinically relevant metrics of plaque vulnerability.

## 1. Introduction

Atherosclerosis is characterized by the thickening of the arterial wall, termed atherosclerotic plaque, at vulnerable sites such as bifurcations and is caused by chronic inflammation (Insull, 2009). Carotid arteries are the main blood supply to the brain. If left untreated, plaques in these vessels can rupture and lead to stroke. Stroke is one the leading causes of death worldwide (Feigin et al., 2021) and carotid atherosclerosis accounts for 34% of all ischemic strokes (Smith-Bindman and Bibbins-Domingo, 2021). The current diagnostic risk criteria in the treatment of patients with carotid atherosclerosis is the stenosis percentage defined by the percentage of luminal narrowing by the plaque build-up. This criterion has proven to be useful for severe carotid stenosis (70-99%) but falls short for mild stenosis patients (0-20%) (Warlow, 1991). Ultrasound (US) imaging is routinely used for carotid plaque assessment because it is fast, relatively cheap, non-ionising, non-invasive and can be easily implemented into a clinical workflow (Liu et al., 2016). More specifically, Doppler US is commonly employed and allows clinicians to infer the percent stenosis in the artery by measuring peak systolic velocity (Grant et al., 2003). Plaques more prone to rupture are typically characterized by the presence of a lipid-rich core and a thin fibrous cap (Saba et al., 2021). Further findings support the argument of plaque composition as a feature of vulnerability. For instance, Schindler et al. showed that the presence of intraplaque haemorrhage is associated with a higher risk of future stroke (Schindler et al., 2020). Recent studies have pointed to the significance of mechanically sensitive indicators to provide pivotal information regarding plaque structural integrity (Ghasemi, Nolan and Lally, 2020). The efforts made in the field ultimately push for a paradigm shift in the risk stratification of carotid plaque vulnerability. To better understand the mechanical behaviour of atherosclerotic plaques, mechanical characterization can be performed ex vivo. Human atherosclerotic plaque tissue is usually harvested from carotid endarterectomy (CEA) surgery. This highly invasive procedure involves either a longitudinal or transverse incision along the neck to reach the carotid artery (Uno et al., 2020). Several studies have carried out uniaxial and bi-axial tensile testing to understand the mechanical properties of intact plaques and components of the plaque (Maher et al., 2009; Teng et al., 2009, 2014; Kural et al., 2012; Johnston, Gaul and Lally, 2021; Tornifoglio et al., 2023). However, these techniques fail to adequately replicate in vivo loading conditions. To counterfeit these limitations, best replicate these conditions and allow for valuable insights into a plaque’s mechanical behaviour, inflation testing can be carried out (Renate W. Boekhoven et al., 2014; Akyildiz et al., 2016; Noble et al., 2020; Guvenir Torun et al., 2021). In these studies, tissue was harvested from cadavers so the arterial wall could be tested as well. Boekhoven et al. carried out inflation testing on intact carotid endarterectomy samples collected from surgeries (Renate W. Boekhoven et al., 2014). The challenge lies in the fact that carotid plaque specimens obtained from CEA procedures are often fragmented or cut longitudinally depending on the surgical approach, rendering pressurization highly challenging if not impossible (Renate W. Boekhoven et al., 2014). To this extent, the use of phantoms as vessel surrogates have many advantages: i) circumventing limited plaque tissue accessibility or availability, ii) enable rupture mechanism testing by offering complete control over material properties and heterogeneity (Sotiriou, Yiannakou and Damianou, 2022), and iii) allowing for validation of computational modelling approaches. One material of particular interest is polyvinyl alcohol (PVA). PVA has been widely used as a tissue mimicking material due to its ability to polymerize and stiffens when subjected to subsequent freeze thaw cycles (FTCs) (Surry et al., 2004). Additionally, it possesses similar acoustic properties to arterial tissue which is of great benefit for studies involving US imaging (Malone et al., 2020). Many studies have sought to investigate the use of PVA to mimic atherosclerotic plaque features (R. W. Boekhoven et al., 2014; Porée et al., 2017; Chee et al., 2018; Gatti et al., 2021). Pazos et al. used intravascular ultrasound (IVUS) to image mock carotid arteries made with a double-layered PVA wall and lipid inclusions (Pazos et al., 2010). Similarly, Widman et al. have characterised hard and soft carotid plaque PVA-based surrogate models through shear wave elastography and mechanical testing (Widman et al., 2015). However, few studies have combined inflation testing of diseased vessel surrogates with non-invasive US imaging and computational modelling. In fact, inverse finite element (iFEM) approaches prove to be a powerful tool to recover material properties in vivo or ex vivo using from different imaging modalities. Many studies in the literature have thus either used stress or strain maps as their ground truth to be matched by their finite element simulations in iFEM approaches (Karimi et al., 2008; Nieuwstadt et al., 2015). Akyildiz et al. developed an iFEM method to determine local material properties of atherosclerotic plaques from ultrasound images used to create 2D finite element models (Akyildiz et al., 2016). Guvenir et al., went further into improving this ultrasound-based framework by introducing a Bayesian optimisation-based pipeline to refine the material property estimations from the iFEM (Guvenir Torun et al., 2021). In this study, we explore the pipeline presented by Narayanan et al., where an iFEM was used to obtain multiple patient specific parameters from optical coherence tomography (OCT) reconstructed atherosclerotic geometries based on an interface matching method (Narayanan et al., 2021). The purpose of this study is to investigate the potential of PVA phantoms as diseased vessel substitutes using a combination of mechanical characterization from ring tensile testing and under pressure loading conditions, US imaging, and computational modelling. Through this study, both experimental and computational approaches can provide invaluable insights into the mechanical behaviour of diseased vessels with the help of vessel surrogates and in turn can help better determine rupture risk from non-invasive ultrasound imaging.

## 2. Materials and Methods

### 2.1 PVA phantom creation

An adaptation of a protocol described in the literature (Malone, 2019) was used where a 10% weight/volume (w/v) PVA solution was created with aluminum oxide (<50nm particle size) as a scatterer. Briefly, PVA powder, aluminum oxide powder, glycerol, and deionized water were mixed and heated to 80-85°C for an hour until a homogeneous solution was obtained. The mixture was then cooled down to 10°C to limit sedimentation of the aluminum oxide particles during the first thermal cycle. Finally, the solution was syringed into 3D printed moulds made from polylactic acid (PLA). The geometric design of these moulds is described in the next section, see Fig. 2. The moulds were maintained at room temperature for an hour to let any air bubbles rise to the surface as any air stuck within the phantom creates artefacts under US imaging. The phantoms then underwent between two and six FTCs. One FTC consists of a controlled freezing phase at -1°C/minute to -15ºC for 14 hours and controlled thawing phase at room temperature for 10 hours.

**Fig. 1.**
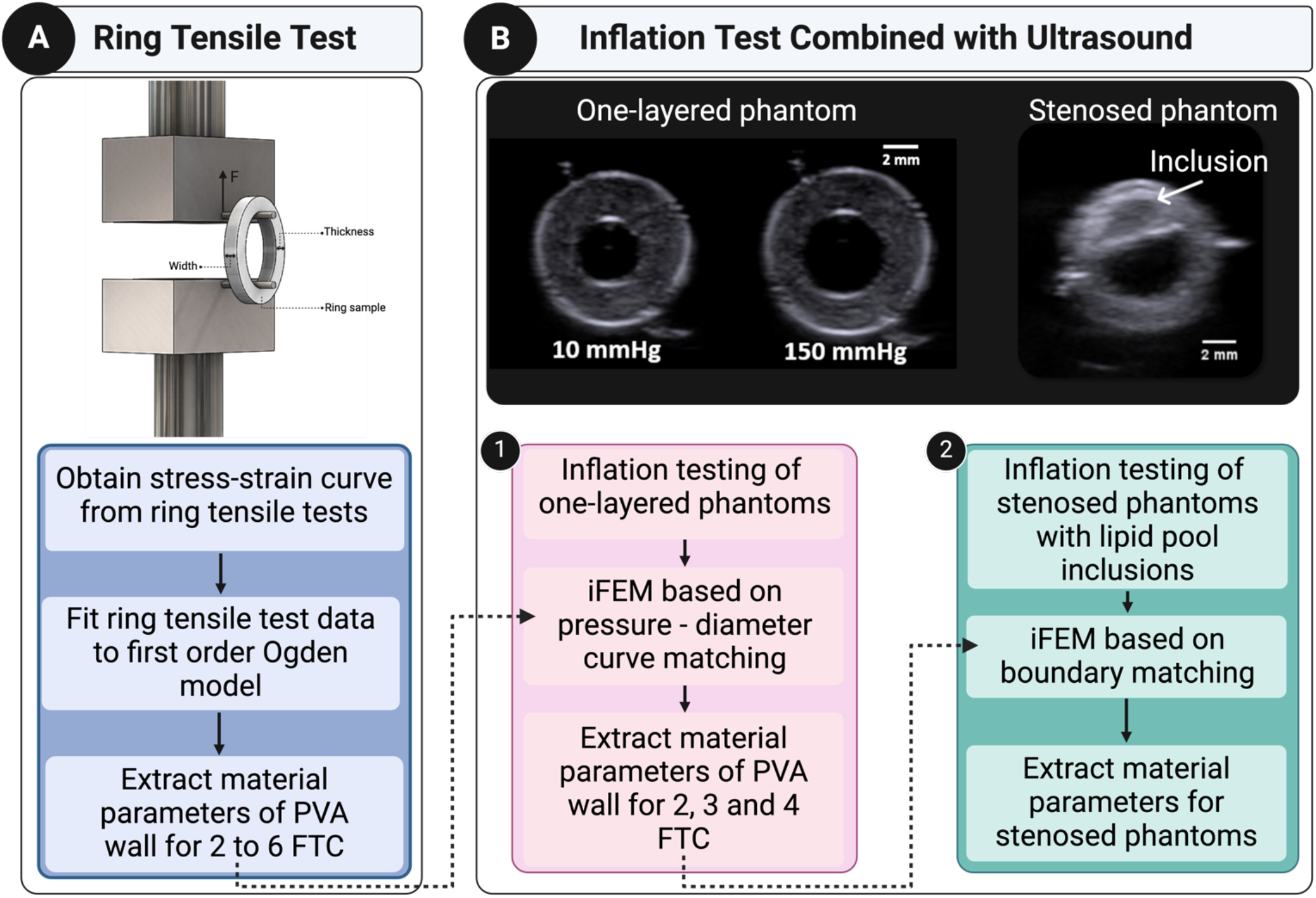
Study overview. Dashed arrows indicate parameters used as initial guesses for the iFEM

**Fig. 2.**
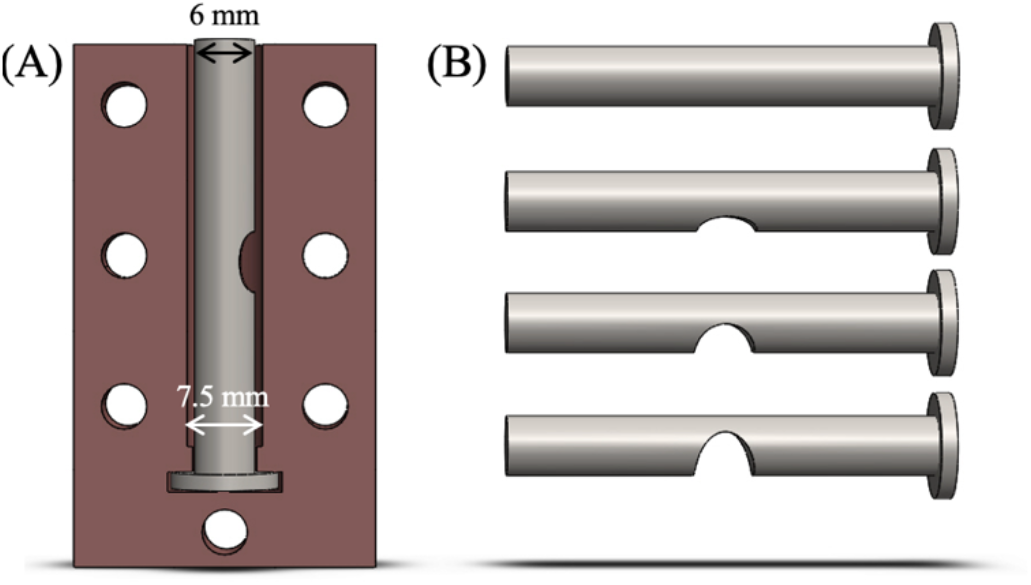
(A) Design of the first mould for bi-layered phantom and (B) inner lumen with 0%, 25%, 50%, 75% stenosis (from top to bottom)

### 2.2 Geometric designs

The phantoms’ moulds were designed using SOLIDWORKS® 3D mechanical CAD software (version 2018, Dassault Systems, S.A., Suresnes, France), see Fig. 2. One-layered phantoms were created using a two-piece hollow cylinder (outer diameter = 9 mm) and a tubular insert (outer diameter = 3.6 mm). To create bi-layer phantoms, moulds were designed using a two-piece hollow cylinder (final outer diameter = 9 mm) and an inserted tube (outer diameter = 6 mm). The empty space was filled with the 10% w/v PVA solution and sealed. A second mould with an inner diameter of 7.5 mm was then placed in the larger diameter mould and another layer injected resulting in two layers each having a wall thickness of 0.75 mm. The bi-layered models underwent physical cross-linking (via FTCs) so that the inner layer would be stiffer (six FTCs) than the outer layer (four FTCs). Fig. 2 shows the core mould design, see Fig. 2 (A) alongside different stenosis percentages on the inserted cylinders, see Fig. 2 (B). Mock vessels with different percent stenoses were created with concave inserted tubes allowing for lumens featuring protrusions of different sizes, 0%, 25%, 50%, 75% stenosis.

### 2.3 Creation of lipid-pool like inclusions

To include a lipid pool-like component within the stenoses, an empty space was created within the wall of the phantom. The empty space was created by placing dissolvable PVA fragments that were printed using an Ultimaker 3 (Ultimaker BV, the Netherlands) in the first layer of the bi-layered phantoms. This layer was placed in water until complete dissolution of the PVA fragments prior to a lipid pool-like substance (Nam *et al*., 2016) being injected into the designated space. The addition of the second layer then covered the first layer as well as the lipid pool inclusion.

### 2.4 Ring tensile test

Ring tensile tests of PVA vessel phantoms were performed for each FTC going from two to six FTCs. The phantoms were made with a nominal mould thickness of 1.0 mm (outer diameter is 7 mm) to have an adequate ratio of the ring thickness to pin radius (Mahutga, Schoephoerster and Barocas, 2021). Mechanical testing was performed at room temperature for each FTC (i.e., two to six FTC with 10% PVA solution). Each tubular PVA was cut into 2-mm rings using a custom 3D-printed microtome blade cutter. The samples were then measured for their thickness and length using a vernier caliper. Three random points of the sample were measured to obtain a mean thickness and length. The samples were submerged in 10% glycerol solution using a transfer pipette before loading on the pins to avoid dehydration. Samples were stretched with a Zwick Roell Z2.5 machine (Zwick Roell Group, Ulm, Germany) with a 20N load cell. The rings were placed in the middle of a custom-made test gauge with a pin radius of 0.5 mm. Samples were set to pre-load of 0.05 N at a speed of 5.0 mm/min. The samples were then pre-conditioned for 5 cycles to 10% strain at a speed of 60% L_o_/min with L_o_ the initial gauge length. The ring samples were stretched until failure with the same speed of the pre-conditioning cycles. The stress-strain curves obtained from the ring tensile test (Macrae, Miller and Doyle, 2016) were used as input to calibrate the material model described in section 2.6 using the Hyperfit software (http://www.hyperfit.wz.cz/). The fitting was performed on a representative sample for each FTC. The mean elastic modulus of each FTC was also obtained.

### 2.5 Combined inflation testing and ultrasound imaging

Fig. 3 illustrates the inflation testing rig developed to incorporate US imaging. US scans were acquired using the SIEMENS ACUSON S2000™ ultrasound system. A 9L4 linear array probe was used with a frequency at 9 MHz to acquire scans. The probe is positioned above the water bath and can be rotated to either obtain cross-sectional or longitudinal views of the phantom. Pressure inflation tests were carried out using a PID controlled PHD Ultra 703007 Syringe Pump (Harvard Apparatus, Holliston, MA, USA) and pressure was monitored with a fiber optic pressure sensor FOP-M260 pressure probe (FISO, Quebec, QC, Canada). The PVA phantoms were axially stretched (10%) and pressurized from 0 to 150 mmHg, in 10 mmHg increments. US B-mode scans were acquired at each pressure step. Overall, three one-layered phantoms (one for each FTC going from two to four) and 12 bi-layered stenosed phantoms (n=3 for each stenosis percentage, 0-75%) were successfully tested and imaged.

**Fig. 3.**
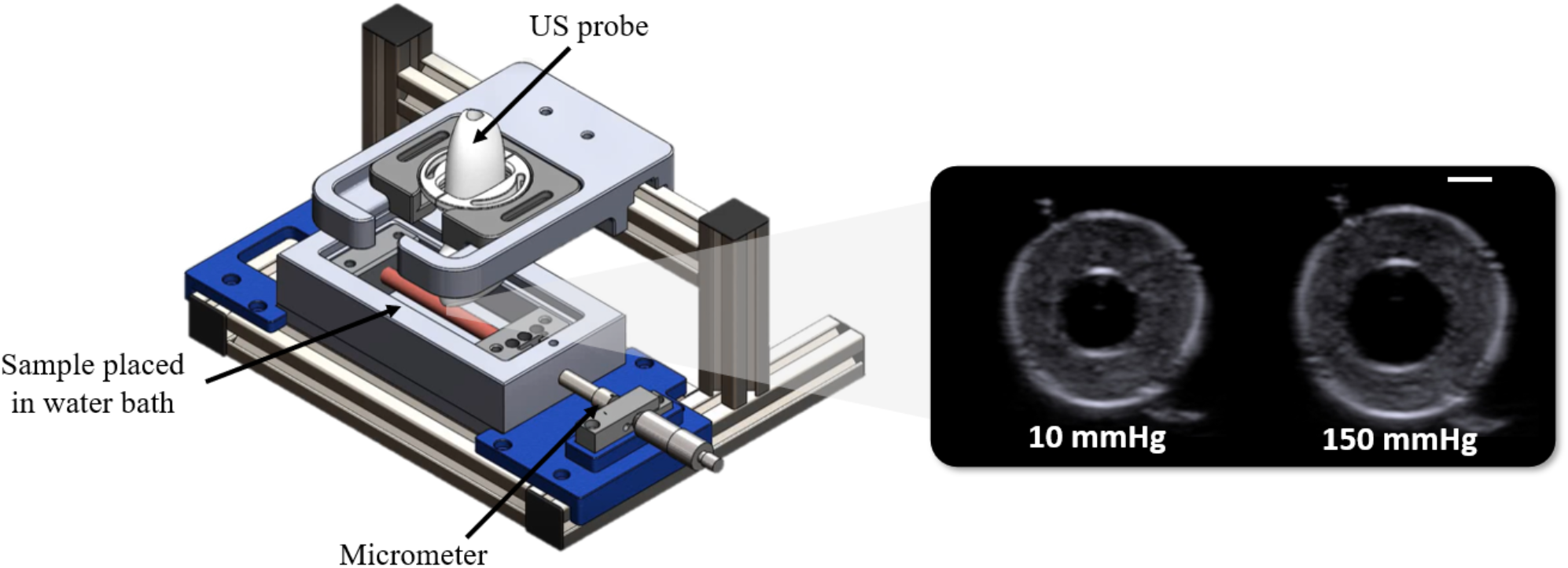
Schematic of inflation testing rig combining non-invasive ultrasound imaging (left) and B-mode scans of one-layered phantom at 10 and 150 mmHg (right). Scale bar: 2 mm.

### 2.6 Constitutive modeling

PVA is a hydrogel that experiences very high deformation and exhibits non-linear stress strain behavior. Several studies have investigated different material models to characterise the mechanical behavior of PVA (Nafo and Al-Mayah, 2020; Fegan *et al*., 2022). In this study, we used a first order Ogden model (stretch-based). The Ogden model is expressed as followed:

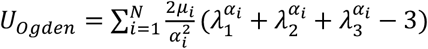

Where *μ* (shear modulus) and *α*_*i*_ (dimensionless) are material parameters and *λ*_1,_*λ*_1_,*λ*_1_ are the principal stretches. For N=1, we thus have:

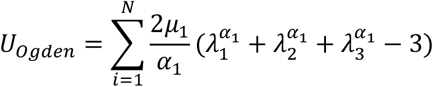

### 2.7 Inverse Finite Element Framework

#### 2.7.1 Diameter matching for one-layered phantoms

Finite element (FE) models of the phantoms were created to model the inflation of the one-layered phantoms (n=3) in Abaqus (Dassault Systemes, 2017) for the 2, 3 and 4 FTC. Hybrid 8-node brick elements with reduced integration (C3D8HR), chosen due to the assumption of material incompressibility, were used to mesh the geometries. The goal of this inverse FE methodology is to determine the optimal material parameters, namely *μ* and *α*, by matching the experimental and computational pressure diameter curves. The objective function *φ* minimised the sum of squared difference (SSD) between the experimental and computational diameters, where the shear modulus *μ* was the optimised parameter, as follows:

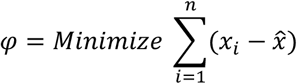

where n is the number of data points, *x*_*i*_ are the observed experimental values and 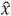 the observed computational values. The optimisation technique followed a Hookes-Jeeves direct search method, and the optimal solution was reached after 50 iterations. This inverse FE methodology was implemented in Isight 5.9 (Dassault Systèmes Simulia corporations, Vèlizy-Villacoublay, France). Briefly, the output coordinates from a created node set on the inner lumen of the phantom model are retrieved from Abaqus. Experimental and computational pressure-diameter curves can then be plotted in the “Data Matching” component in which the comparison metric, the sum of squared difference (SSD) is defined and used in our optimisation procedure.

#### 2.7.2 Diameter matching for bi-layered phantoms

After data acquisition, an open-source toolbox, namely Ultrasound Research Interface Offline Processing Tools (URI-OPT), was used to read in the beamformed RF data as a matrix for each frame acquired (Brunke *et al*., 2007). The outer, inner lumen and lipid pool were segmented in a two-step process using an in-house MATLAB (The Mathworks Inc., Natick, MA) code. Firstly, a freehand tool was used to define the initial region of interest for each delineation. The latter was then used as an input for an active contour segmentation algorithm. Due to the inherent scattered nature of the ultrasound images, the outlines output from the active contour function were irregular. A smoothing step was thus needed and was performed by importing the boundary coordinates in SOLIDWORKS® (version 2018, Dassault Systems, S.A., Suresnes, France). Using the curve wizard tool, the contours were smoothed and exported as an .igs file. A sketch was then created in Abaqus (Dassault Systemes, 2017) to define a 2D cross-section of the PVA phantom. The nodal coordinates of the boundaries of interest (i.e., inner, outer lumen and lipid pool) were exported in an excel file and read into a MATLAB (The Mathworks Inc., Natick, MA) code to obtain the final smoothed region of interest. For more complex cross-sectional geometries, a boundary matching inverse approach was proposed following the framework presented in (Narayanan *et al*., 2021). Briefly, if we name the target geometry ℊ_*target*_, at intraluminal pressure *P*_*final*_and the optimised geometry ℊ_*Opt*_ with an intraluminal pressure *ΔP* = *P*_*final*_–*P*_*intial*_, the objective was to determine optimized parameters that will allow for the boundaries of ℊ_*Opt*_ to match the ones of the target geometry ℊ_*target*_. As shown in Fig. 4, the aim was to match boundaries of the optimised pressurised geometry to the ground truth geometry contours. We modelled the 2D cross sections of the phantom’s wall as first order Ogden models where *μ*_*wall*_, *α*_*wall*_ are the parameters to be optimised. Initial material parameters of the wall were taken from the one layered 4 FTC phantom optimized parameters, see Fig. 4. Additionally, the lipid pool was defined as a Neo-Hookean model with an initial *C*_10_ of 1 kPa (Guvenir Torun *et al*., 2021). Plane strain hybrid elements (CPE4H) were used for the finite element simulations. An inverse FE algorithm was implemented to adjust the parameters to the inflation testing data using Isight 5.9 (Dassault Systèmes Simulia corporations, Vèlizy-Villacoublay, France). A Python code was developed to calculate the sum of the minimal Euclidean distance between the nodal coordinates of the respective boundaries of the optimised 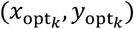 and target geometry 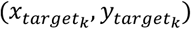. The final error *ϵ* to be minimized was defined as follows:

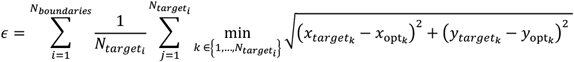

where *N*_*boundaries*_ is the number of boundaries to be matched, 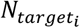 is the number of nodes in the target geometry of the interface *i*, and 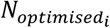 is the number of nodes on the optimised geometry of the interface *i*. Fig. 4 shows the iSight workflow. Briefly, an input file is read into the “Abaqus component” where the material parameters to be optimized are selected. Material parameters from the inflation test calibration were used as initial guesses. The in-house Python script was run in Abaqus where the .odb file was taken as the input to calculate the error and write it to a text file. This text file was then read in the “Data Exchanger” component to create the error variable. Finally, the error minimization was carried out in the “Optimisation” component where a Hookes-Jeeves algorithm was used to identify optimal parameters.

**Fig. 4.**
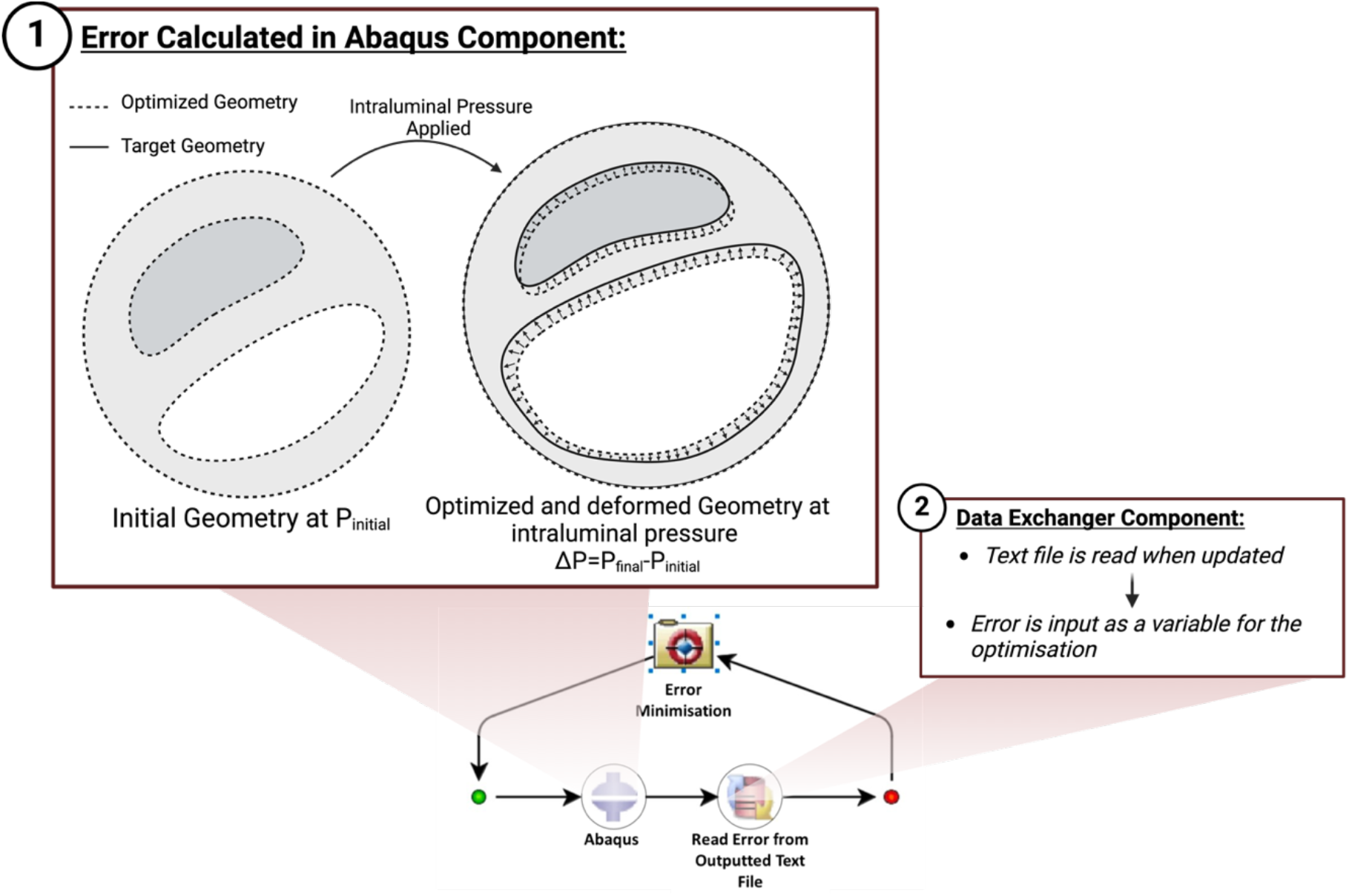
iSight framework schematic.

### 2.8 Statistical analysis

Statistical analysis was performed with Prism 6 statistical software (GraphPad Software Inc., San Diego, California). All data was tested for normality using Shapiro-Wilk normality tests and equality of group variances using Brown-Forsythe ANOVAs. All data in this study passed normality tests. In the case of unequal group variances, Brown-Forsythe and Welch ANOVAs with Dunnet’s T3 multiple comparison tests were used, otherwise ordinary one-way ANOVAs with Tukey’s multiple comparison were used. For the pressure-diameter experimental data, a one-way ANOVA was performed to investigate statistical significance (i.e., p < 0.05) between each of the tested stenosed phantom groups (0%, 25%, 50%, 75%).

## 3. Results

### 3.1 Ring tensile test

Stress strain curves from the ring tensile tests of each FTC group show strain stiffening behavior typical of arterial tissue, specifically when the number of FTC increases, see Fig. 5 (A). Looking at the mechanical data of all 35 rings tested between 2 and 6 FTC, stiffening and strengthening is observed with increased numbers of FTCs. This is further shown in Fig. 5 (B) where the ultimate tensile (UT) stress of the 2 FTC group (0.14 ± 0.03 MPa) is significantly lower than the 3 FTC (0.33 ± 0.03 MPa) and 6 FTC (0.86 ± 0.27 MPa) groups. Groups 4 FTC (0.49 ± 0.32 MPa) and 5 FTC (0.49 ± 0.32 MPa) did not show any significant difference compared to the other groups. Fig. 5 (C). shows a similar trend where an increase in the number of FTCs leads to higher UT strain values. The 2 FTC group had a significantly lower UT strain (0.44 ± 0.09) compared to the 3 FTC (0.68 ± 0.09), 5 FTC (0.77 ± 0.22) and 6 FTC (0.93 ± 0.18) groups. No statistically significant difference between the 4 FTC (0.76 ± 0.34) group and the other sample groupings was observed, however. Table 1 shows the average elastic modulus of the 2, 3, 4, 5, 6 FTCs to be equal to 331 ± 21, 510 ± 43, 630 ± 110, 662 ± 110, 957 ± 147 kPa, respectively. Fig. 6 illustrates the fitted 1-term-Ogden model to the experimental data of a representative sample for 4 FTCs with high goodness of fit. The results of the fitted models for each FTC are depicted in more detail in Table 2 with a shear modulus *μ*_1_ ranging from 90 kPa to 235 kPa respectively for 2 to 6 FTC. The *α*_1_ value did not vary much between FTCs ranging from 4 to 4.8.

**Table 1.** Elastic Modulus (Mean ±SD) of each FTC.

| Mean elastic modulus |  |  |  |  |  |
| --- | --- | --- | --- | --- | --- |
|  | 2 FTC | 3 FTC | 4 FTC | 5 FTC | 6 FTC |
| $E$ (kPa) ± SD | $331 \pm 21$ | $510 \pm 43$ | $630 \pm 110$ | $662 \pm 110$ | $957 \pm 147$ |

**Table 2.**
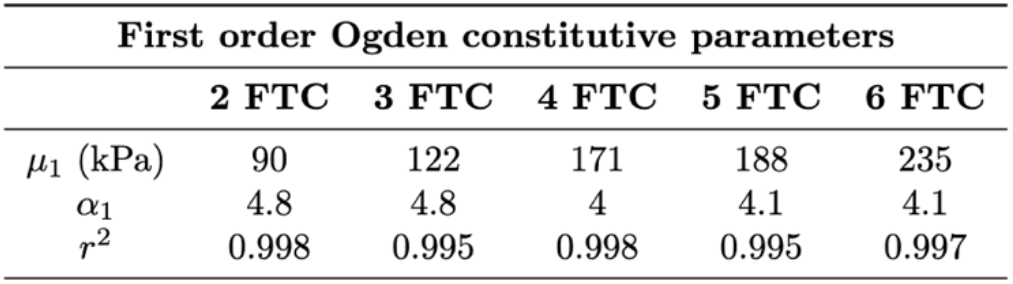
Estimated material parameters of the PVA ring tests for the first order Ogden constitutive models with the coefficient of determination (R^2^) for each sample.

**Fig. 5.**
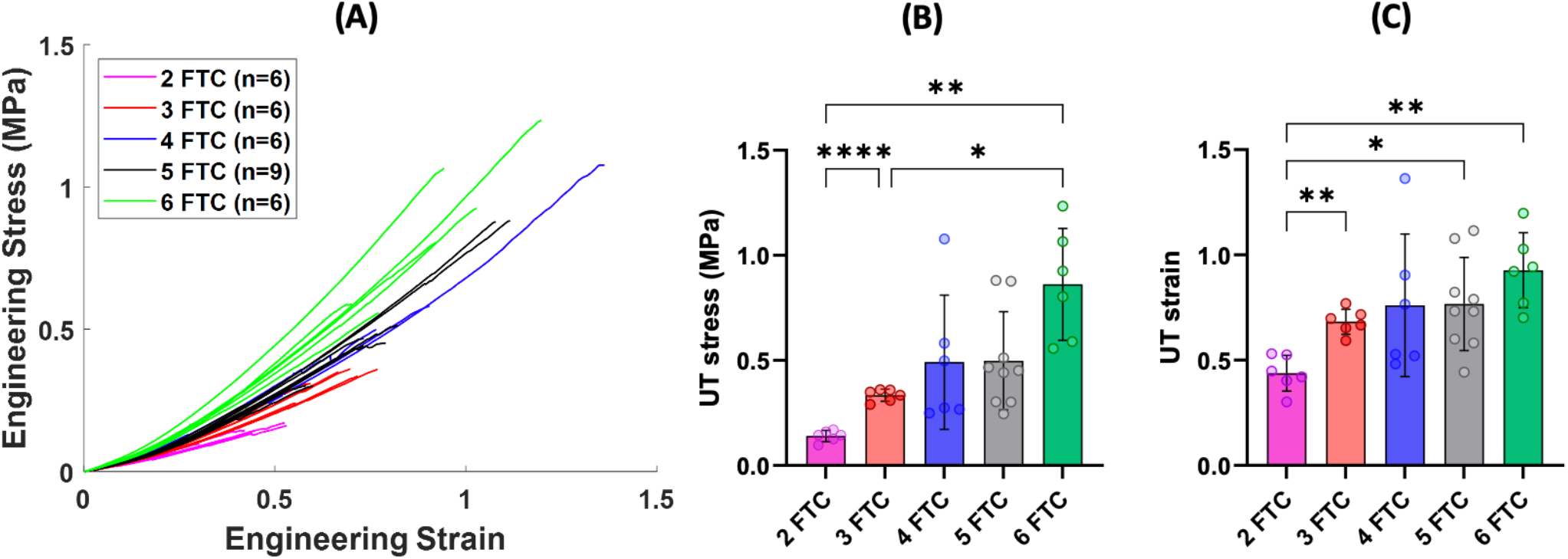
(A) Stress-strain curves for n=35 rings, color coded by their respective groupings. (B) UT stress: significance determined by Brown-Forsythe and Welch ANOVA with Dunnett’s T3 post hoc multiple comparisons, 2 FTC and 3 FTC ****p<0.0001, 2 FTC and 6 FTC **p=0.0083, 3 FTC and 6 FTC *p=0.0317. (C) UT strain: significance determined by Brown-Forsythe and Welch ANOVA with Dunnett’s T3 post hoc multiple comparisons, 2 FTC and 3 FTC **p=0.0001, 2 FTC and 5 FTC *p=0.0177, 2 FTC and 6 FTC **p=0.0042.

**Fig. 6.**
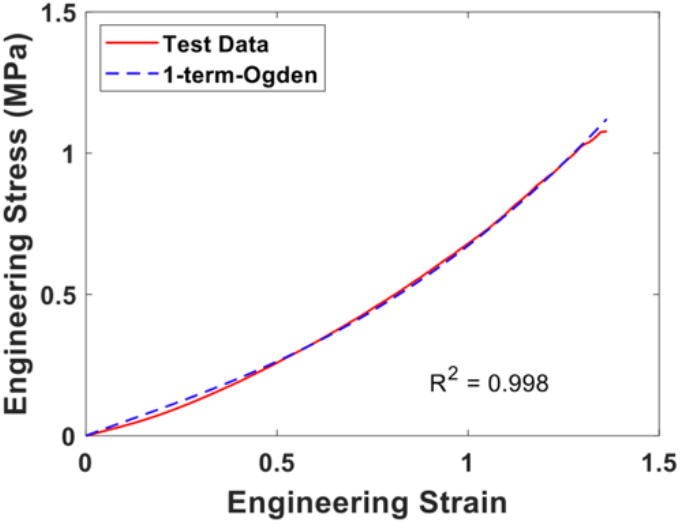
First order Ogden model fitted to the stress-strain curve of a 4 FTC sample.

### 3.2 Inflation of bi-layered phantoms

Bi-layered stenosed phantoms featuring lipid-pool-like inclusions were successfully inflated and imaged using US from 0 to 150 mmHg, see Fig. 7. The hypoechoic (dark grey areas indicated by the white arrows in Fig. 7) reveal the presence of the lipid-like content in the occluded regions. The red dashed circles indicate regions known as the plaque “shoulders”. It is interesting to qualitatively note that a higher displacement was observed at the shoulders to the rest of the phantom wall, further quantified in Fig. 11.

**Fig. 7.**
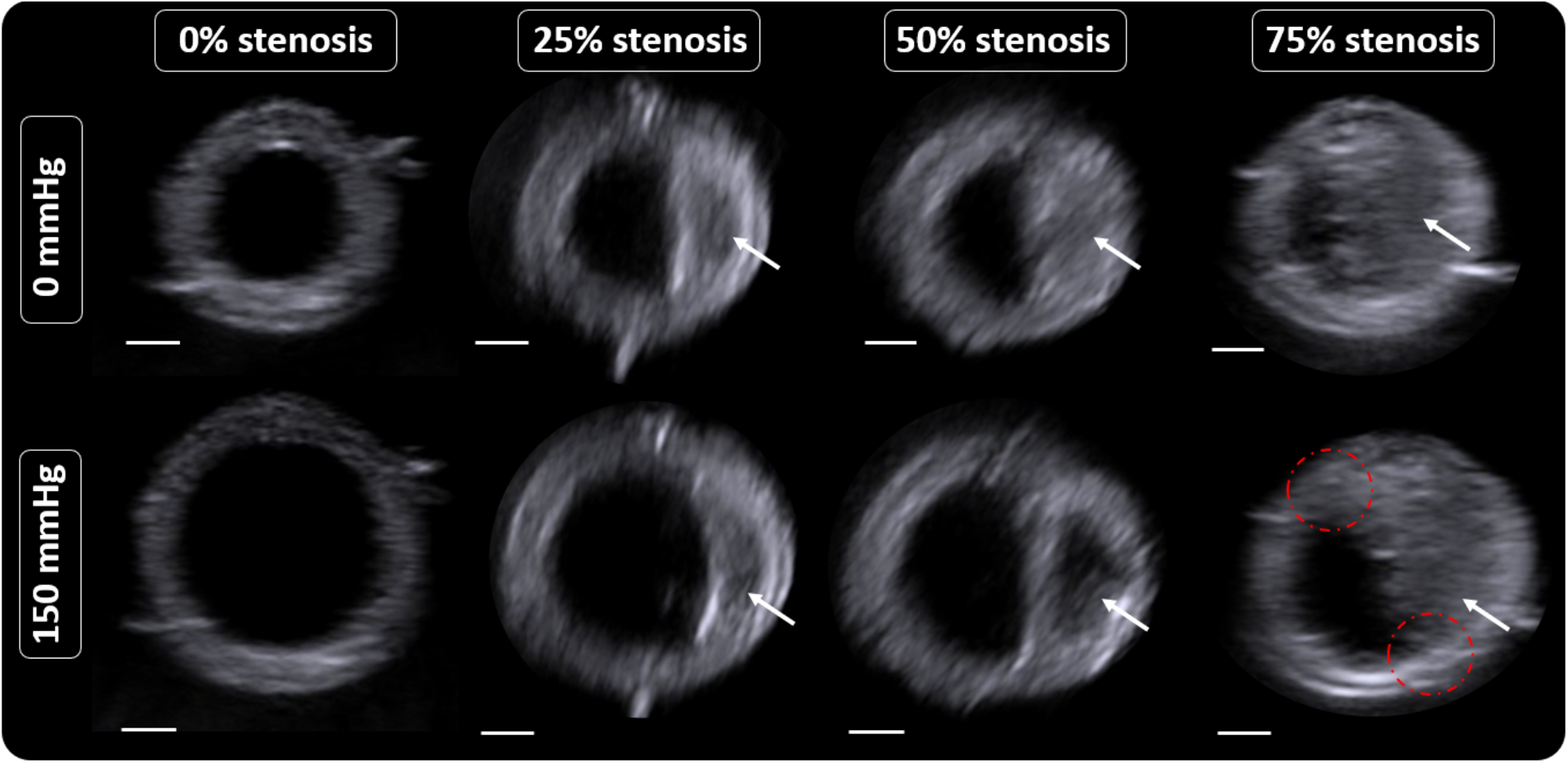
B-mode ultrasound of PVA phantoms. Scale bars: 2 mm

Lumen area is plotted as a function of pressure in Fig. 8 (A) as it provides a better representation of the lumen’s elliptical shape compared to a single diameter measurement. The solid lines in Fig. 8 (B) represent the mean from n=3 samples at each stenosis level, while the shaded areas represent the standard deviation. Additionally, the slope of the lumen area versus pressure curve (across the total pressure range of 150 mmHg) is given in Fig. 8 (B). The latter shows that the area expansion of the 0% (2.4 ± 0.53 cm^2^\mmHg x 10^−3^) stenosis samples is significantly higher than that of the 25% (1.4 ± 0.14 cm^2^\mmHg x 10^−3^), 50% (1.2 ± 0.08 cm^2^\mmHg x 10^−3^) and 75% (0.73 ± 0.24 cm^2^\mmHg x 10^−3^).

**Fig. 8.**
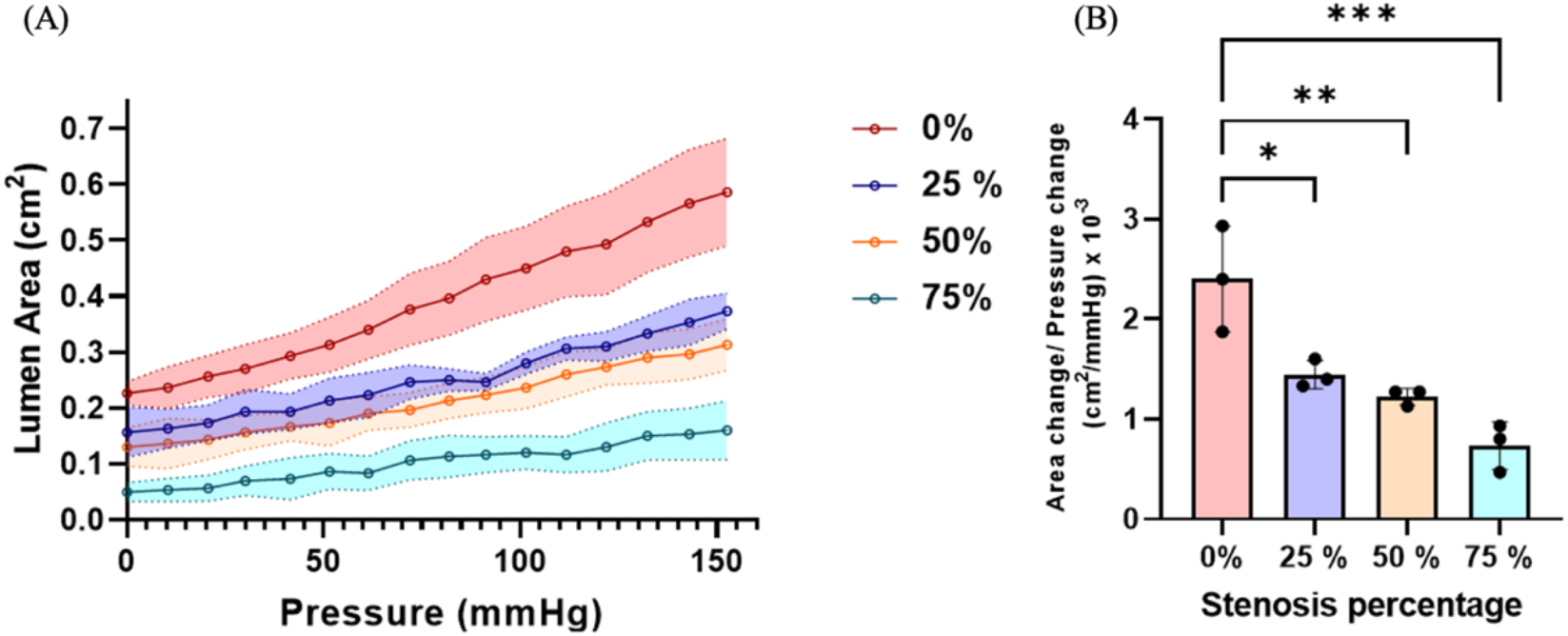
(A) Lumen area against pressure curves for 0%, 25%, 50% and 75% groups and (B) bar plot of the slope of the curves in (A) for each specimen group. Significance determined by multiple comparisons one-way ANOVA with Tukey’s multiple comparisons test; *p=<0.05, **p=<0.01 and ***p=<0.001.

### 3.3 Inverse FE

#### 3.3.1 Diameter matching for one-layered phantoms

Experimental pressure diameter curves did not exhibit strain stiffening behaviour but rather a near linear relationship. Optimised material properties of homogeneous PVA phantoms were recovered by matching the experimental and computational pressure diameter curves of 2, 3 and 4 FTC one-layered phantoms as illustrated in Fig. 9. The latter shows good matching of the computational results with the experimental pressure diameter curves, most notably as the sample gets stiffer – i.e., increasing number of FTCs. For the 2, 3, 4 FTCs, the shear moduli and alpha values recovered were respectively *μ*_1_= 40 kPa and *α*_1_= 6.7, *μ*_1_= 82 kPa and *α*_1_= 9.7, *μ*_1_= 191 kPa and *α*_1_= 9.2, see Fig. 9.

**Fig. 9.**
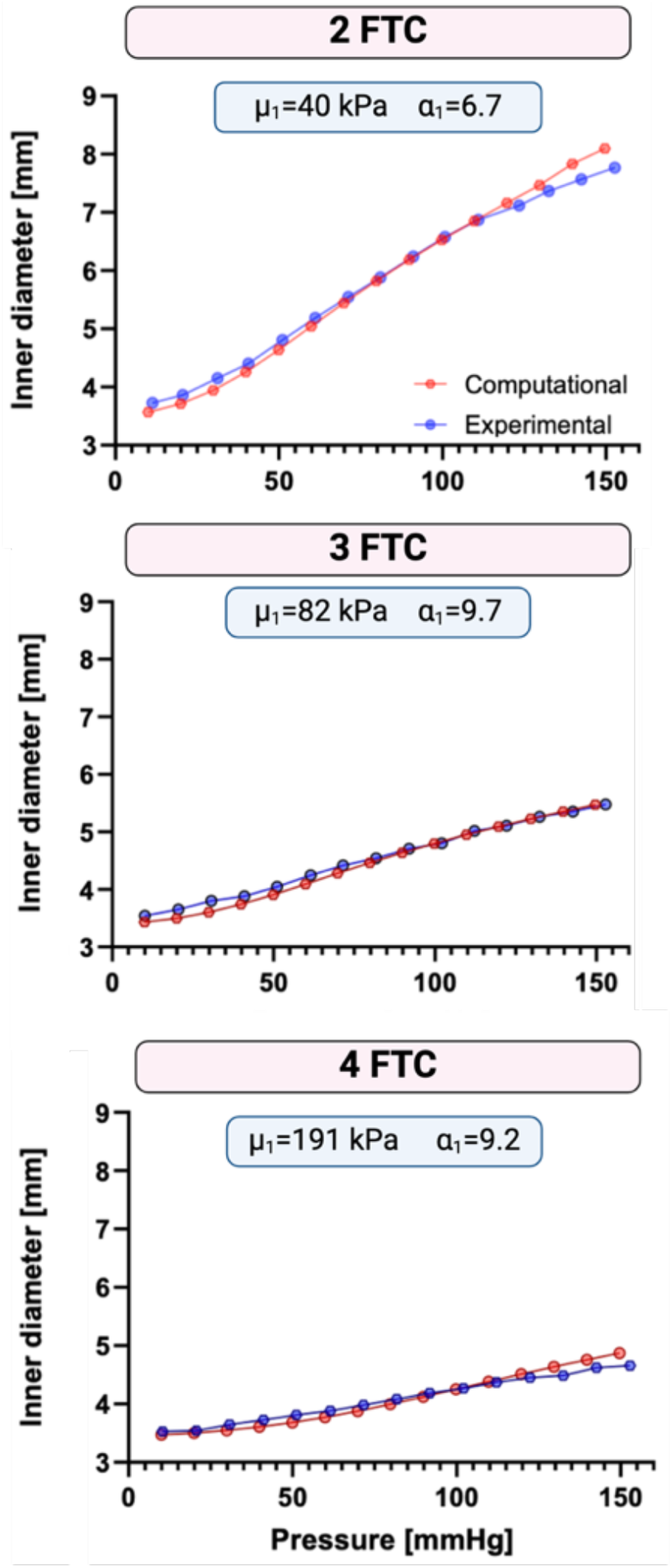
Experimental and computational pressure diameter curves for 2, 3 and 4 FTC with optimal first order Ogden parameters.

#### 3.3.2 Boundary matching

The proposed boundary matching framework, based on the approach by (Narayanan *et al*., 2021), was used to estimate *μ*_1*wall*_ and *α*_1*wall*_ for the PVA wall and the *C*_10_ parameter for the lipid-pool from the inflation testing data between 80 and 120 mmHg for stenosed phantoms with lipid pool-like inclusions. The final recovered shear moduli for the PVA ranged between 27 and 53 kPa. In the cases where the lipid was included (i.e., 25% and 50%), the recovered *C*_10_ values were respectively equal to 0.1 kPa and 0.38 kPa. The latter values are within the range of lipid pool material properties chosen in the literature for inverse FE modelling (Guvenir Torun *et al*., 2021). The final error *ϵ* defined in Section 2.a was minimized for n=1 sample of each stenosis percentage group i.e., 0%, 25%, 50% and 75%. It is important to note that the lipid-pool could not be delineated for the 75% sample hence only two boundaries were matched, namely the inner and outer lumen boundaries. Fig. 10 illustrates the final minimized error and recovered material parameters of the presented inverse FE methodology for each percent stenosis group. The 0% stenosis sample, with two boundaries to match, had the lowest final error with a value of 0.28 mm and a percentage of area difference of 1.4%. On the other hand, the highest error, 0.5 mm (5.7 % percentage of area difference), was reached for the 50% stenosis sample with three boundaries matched including the lipid pool. The 25% and 75% stenosis phantoms, respectively, reached errors of 0.48 mm (2.5% percentage of area difference) and 0.45 mm (2.7% percentage of area difference). Maximum principal engineering strain and Von Mises stress contour plots are shown in Fig. 11 for each stenosis percentage. The presence of the lipid core yields higher strains, see Fig. 11. This is specifically notable for the 75% stenosis phantom, where a maximum principal in-plane engineering strain of 0.33 is present at the shoulder regions. A maximum Von Mises stress was also observed for the 75% stenosis, at the shoulder region, with a peak value of 34 kPa.

**Fig. 10.**
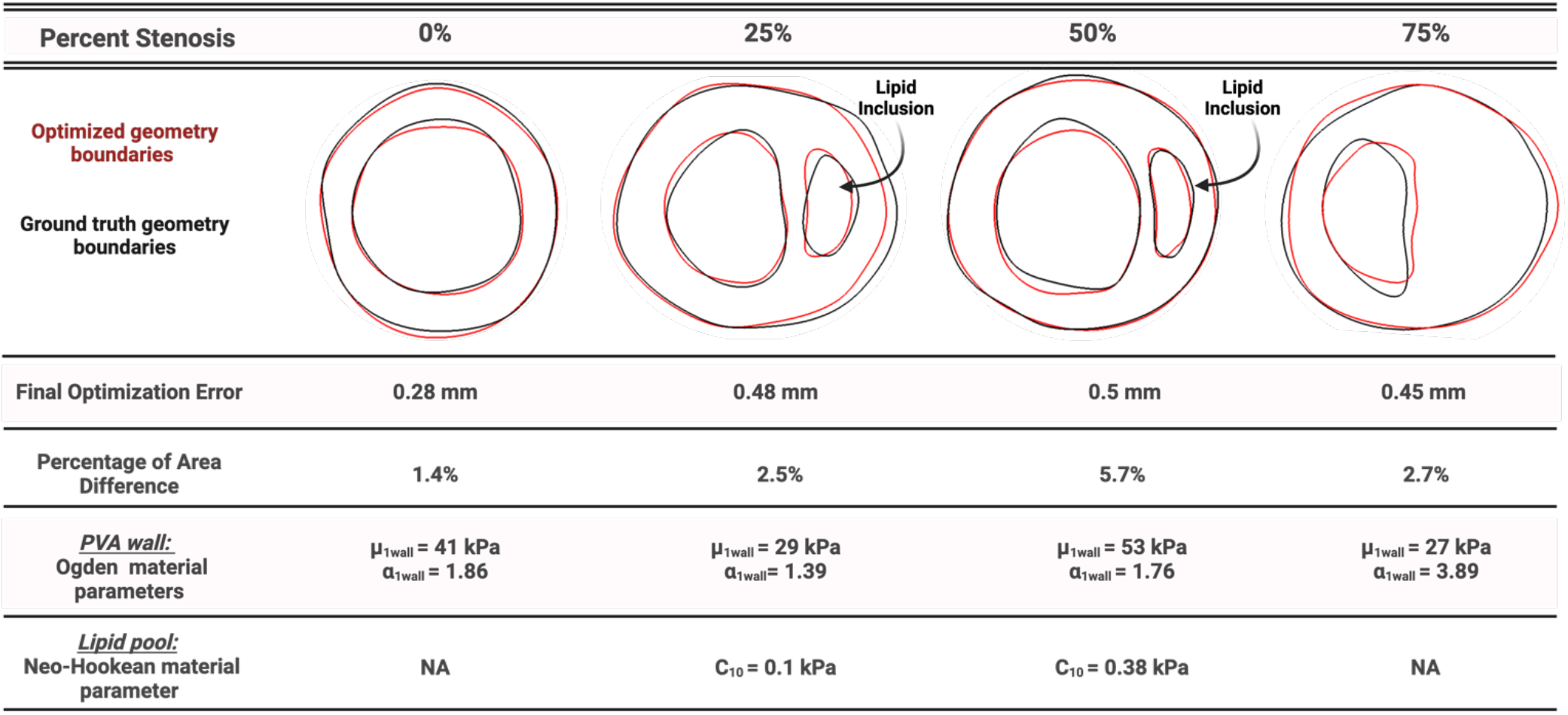
From top to bottom, contours visualisation of the target (red) and deformed (black) geometries, final errors of the optimisation process, percentage of area differences, PVA wall Ogden optimised material parameters and lipid pool optimised parameters for the 25% and 50% stenosis.

**Fig. 11.**
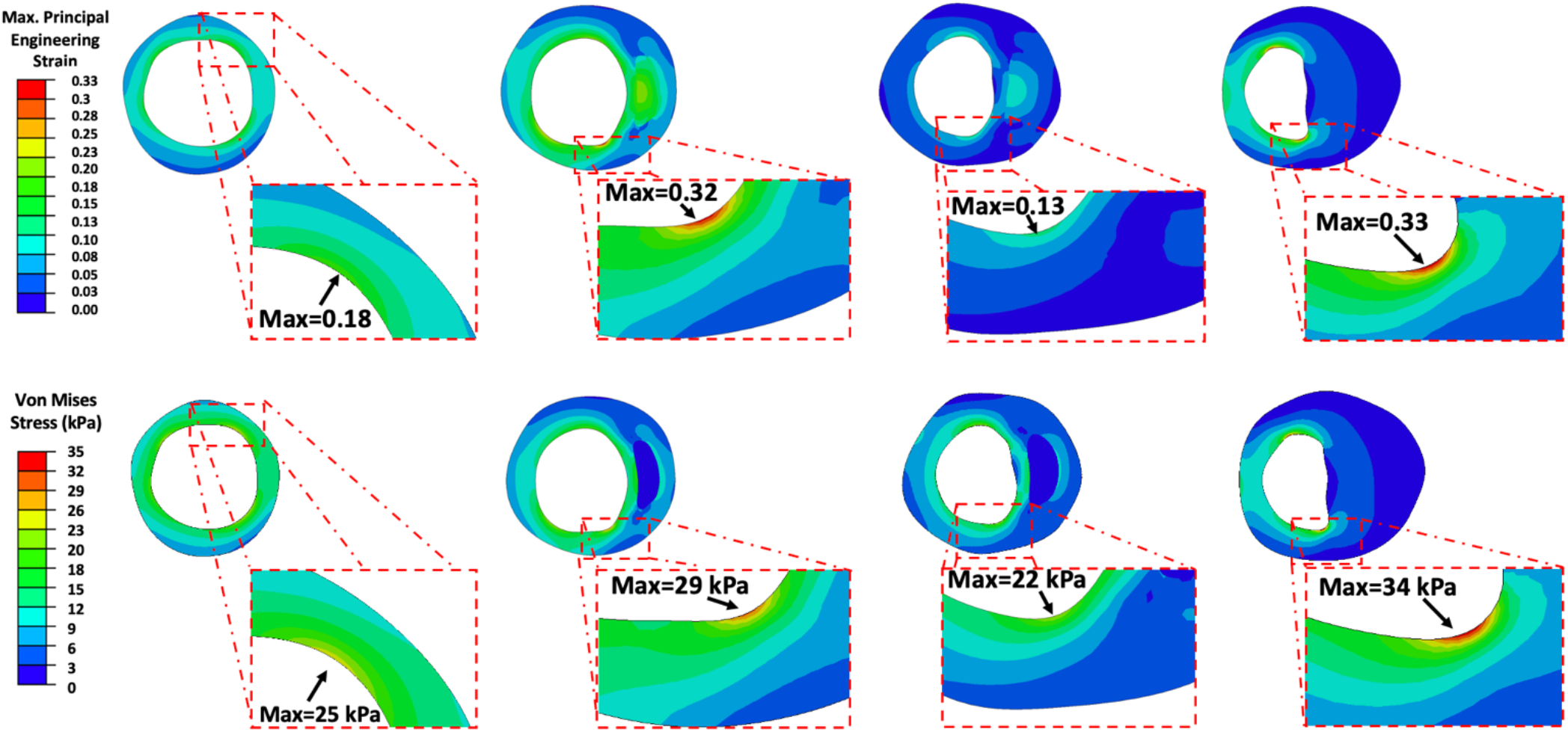
Maximum principal engineering strain (top) and Von Mises stress (bottom) contour plots for 0%, 25%, 50% and 75% stenosis (left to right).

## 4. Discussion

Through a combination of experimental and computational methodologies, this study highlights the potential of PVA phantoms to offer key insights into diseased vessel mechanical behaviors. As illustrated through the results from the ring tensile testing, PVA has tunable material properties; specifically, it stiffens with an increasing number of FTCs with shear moduli ranging between 90 and 235 kPa. In fact, this behavior has been previously shown in several studies illustrating the cross-linking phenomenon induced by the freeze-thaw process (Fromageau *et al*., 2003; King *et al*., 2011; Malone *et al*., 2020). King et al. performed uniaxial tensile testing and found that the stiffness of 10% wt PVA specimens reaches a plateau after five FTCs with Young’s modulus values between 19 and 165 kPa (King *et al*., 2011). Comparatively, the averaged values of elastic modulus from the ring test ranged between 331±21 kPa and 957±147 kPa exceeding the higher range documented in the King et al. A similar levelling trend starting from 4 FTCs was found in Malone *et al*. who carried out uniaxial tensile tests on dog-bone and vessel shaped tubular samples (Malone *et al*., 2020). The stress strain curves from the ring tensile test exhibit a hyperelastic behavior especially for higher FTCs. Khamdaeng et al. assessed carotid stiffness *in vivo* using an ultrasound based technique in seven healthy volunteers (Khamdaeng *et al*., 2011). The reported average moduli before and after the transition point (slope change in stress strain curve) were equal to 160±40 kPa and 900±250 kPa. Franquet et al., evaluated the elastic modulus of the common carotid *in vivo* using MRI and found a 38% increase between the ‘Young healthy subjects” (YH) ‘Old diseased patients’(OD) with respective values of 315±29 kPa and 509±65 kPa (Franquet *et al*., 2013). These values overlap with the elastic moduli found in our study, highlighting the potential for PVA to mimic arterial tissue properties.

A comprehensive study by Fegan *et al*. reported first order Ogden and Yeoh constitutive material coefficients for 10% wt PVA/gelatin samples fitted to compression tests (Fegan *et al*., 2022). The results in Fegan et al. agree with our findings; revealing that the first Ogden model provided a good fit for the PVA phantoms’ stress strain curves. Moreover, the shear moduli gathered from the ring tensile testing and inflation testing in our study are on a higher range compared to those found in Fegan et al., 2022 with values ranging between 22 to 44 kPa. It should be noted that this discrepancy might be due to the fact that values we compare are extracted from different types of mechanical tests (Fegan *et al*., 2022).

Additionally, the shear moduli extracted from the inflation testing of one-layered phantoms in our study sit well within the range of porcine native tissue. In fact, studies on porcine carotid arteries, using both inflation testing and bi-axial tensile testing, have estimated the shear modulus of a porcine carotid arteries to range between 48 – 65 kPa (Vychytil *et al*., 2010; Boekhoven *et al*., 2016); highlighting that PVA subjected to two and three FTCs is a mechanically-suitable healthy vessel surrogate.

PVA also holds significant promise when it comes to replicating diseased and atherosclerotic geometries (Malone et al. 2020). Tornifoglio et al., 2023 unaxially tested human atherosclerotic plaques yielding a mean elastic modulus of 1.26±0.6 MPa (Tornifoglio *et al*., 2023). Plaque caps tested uniaxially by Johnston at al., 2021 exhibited a mean stiffness of 0.05±0.03 MPa in the axial direction versus 0.19±0.07 MPa in the circumferential direction (Johnston, Gaul and Lally, 2021). Davis et al. performed tensile testing on fibrous caps and determined a high strain modulus of 1029.8±795.4 kPa (Davis *et al*., 2016). The elastic modulus values identified in this study align with those reported in existing literature for plaque tissue. However, it is important to note the considerable mechanical variability inherent in such tissue. Therefore, PVA emerges as an optimal material choice for replicating this broad mechanical range via varied concentrations and FTCs (Malone *et al*., 2020).

Previous studies have investigated the creation of PVA phantoms with varying degrees of stenosis (Chee *et al*., 2018; Gatti *et al*., 2021), lipid pool inclusions (Boekhoven et al., 2014; Porée et al., 2017), and hard inclusions (Porée *et al*., 2017). Le Floc’h et al. were the first to create a vessel with a necrotic core or soft inclusion, made from PVA subjected to one FTC (Le Floc’h *et al*., 2010). The versatility of PVA also permits the creation of complex geometries to better imitate specific anatomies such as the carotid bifurcation (Chayer *et al*., 2019). Many of these studies have looked into the use of those phantoms for flow evaluation and stiffness measurements; however, few focus on their mechanical behaviour (Boekhoven,et al., 2014; Nafo & Al-Mayah, 2020).While uniaxial tests have been performed to provide understanding of the mechanical properties of PVA (Surry *et al*., 2004; Nafo and Al-Mayah, 2020), these do not accurately reproduce the multi-axial nature of physiological loadings. Nafo and Al-Mayah carried out uniaxial tests and cavity expansion (pressure with equi-biaxial tension) tests of PVA hydrogel specimens that underwent two FTCs (Nafo and Al-Mayah, 2020). Three models were used to predict the PVA shear moduli, namely, the Ogden, Yeoh and Arruda-Boyce models. Nafo and Al-Mayah reported shear moduli ranging between 20 and 26 kPa, values that are below what we obtained for 2 FTC from ring tensile test (90 kPa) and inflation testing (40 kPa).

This work expands on these previous studies by investigating bi-layered phantoms featuring different stenoses and lipid pool inclusions. Fig. 7 illustrates the characteristic speckle pattern of arterial tissue successfully reproduced by the PVA phantoms. While differentiating the two layers was not possible, we were able to delineate the lipid inclusion – hypoechoic regions – for the 25% and 50% stenosis samples. Moreover, Fig. 7 highlights the uniform wall expansion of a 0% stenosis bi-layered phantom, a result very characteristic of a healthy artery. On the other hand, the 75% stenosed sample in Fig. 7, shows an inhomogeneous expansion of the vessel wall – with a higher displacement in the “shoulder” regions circled in red. And further displayed in Fig. 11 with a peak engineering strain of 0.33 in the 75% stenosis sample. This aligns with findings from (R. W. Boekhoven *et al*., 2014) who used inflation testing of PVA phantoms mimicking fatty plaques and observed higher strains collocated with the fatty pool, which we can visualize as well for the 25, 50 and 75% stenosis samples in Fig. 7. These observations show great promise and warrant further investigations into the development of surrogates to decipher the mechanical behavior of diseased tissue. Additionally, pressure-lumen area curves in Fig. 8 (A) showed that higher stenosis percentage yields decreased area expansion. This was further revealed in Fig. 8 (B), where the slope of the curve for the 0% stenosed specimen (2.4 ± 0.53 cm^2^\mmHg x 10^−3^) was significantly higher than for all other stenosed groups across the same pressure range. This corroborates with findings in C.P Loizou et al. who found a linear relationship between the percent stenosis in the internal carotid artery and the percent of carotid distensibility with a decrease of distensibility when the stenosis percentage increases (Loizou *et al*., 2021).

To further characterize the stenosed vessel surrogates, this study presents a boundary matching based inverse FE framework. This approach demonstrates very good agreement between the boundaries delineated from the inflation test (between 80 and 120 mmHg) and the FE simulations with the highest percentage of area difference of 5.7% for the 50% stenosis phantom, see Fig. 10. Through the combined use of inflation testing and US imaging, insightful information on the behavior of these bi-layered stenosed models was thus revealed. It was seen that the highest stenosis percentage leads to the highest strain and stresses at the shoulder regions, Fig. 11. This aligns with findings in the literature (Li *et al*., 2006; Sadat *et al*., 2010; Ghasemi, Nolan and Lally, 2020). The stress ranges found in Li et at al., were however on a higher scale than in this study ranging from 161.1 kPa to 694.1 kPa. Soleimani et al. also found that the mean effective, circumferential and first principal stress values increased when the stenosis increased from mild to significant in carotid arteries (Soleimani *et al*., 2021).

Some limitations need to be acknowledged in this study. Firstly, the use of non-invasive ultrasound imaging to delineate material boundaries can prove to be non-trivial. In fact, for the 75% stenosis phantom, manual delineation of the lipid inclusions and wall layers was not possible. The PVA wall was thus considered as a whole for the inverse FE framework with only one shear modulus value to be optimised. In Narayanan et al., a deep learning method for OCT images was used to classify the microstructural components in five classes: calcium, mixed, lipid, fibrous and healthy wall (Narayanan *et al*., 2021). This limitation can however be overcome using ultrasound-based imaging techniques such as elastography for instance. Hence, Le Floc’h et al. developed an IVUS based elastography map to accurately detect soft inclusions within PVA phantoms and were able to characterise their Young’s moduli (Le Floc’h *et al*., 2010). Furthermore, ultrasound machines such as the Aixplorer were specifically designed to quantify tissue elasticity of soft and hard regions using shear wave elastography. Widman et al. constructed n=6 PVA based carotid phantoms with soft and hard inclusions (Widman *et al*., 2015). An Aixplorer ultrasound machine was used to acquire shear wave elastography (SWE) measurements of the phantoms. The shear moduli yielded from SWE for the soft and hard plaques were respectively 5.8 ± 0.3 kPa and 106.2 ± 17.2 kPa. Strain based methods have also proven to be efficient to delineate components. More recently, Latorre et al. introduced a strain gradient-based plaque segmentation framework (Latorre et al., 2022). This approach was able to delineate the fibrous cap thickness and lipid pool from simulated data with a Segmentation Index (SI) related to the segmentation performance, above 90%. An expanding field, namely photoacoustic imaging, has also shown promising results concerning the discrimination of plaque components. In this regard, Cano et al. used multispectral photoacoustic imaging on plaque PVA phantoms and plaque tissue to accurately decipher plaque components (Cano *et al*., 2023).

Additionally, we did not incorporate anisotropy in our study. A non-linear stress-strain curve would be expected in inflation testing of arteries due to their anisotropy (Boekhoven *et al*., 2016) whereas PVA is more homogenous and shows a near linear behaviour in our inflation testing. In future work, this phenomenon will be investigated through the addition of fibres within the PVA wall. In fact, melt electrowriting can be used to design fiber arranged patterns to mimic native tissue architecture (Federici *et al*., 2023). Inducing anisotropy in PVA could also be achieved by stretching the PVA hydrogel while it is being cross-linked (Chatelin *et al*., 2014). Bernal et al. developed anisotropic aortic models and evaluated their shear modulus using a custom-made ultrasound probe (Bernal *et al*., 2019). They observed an increase in the shear modulus as a function of pressure from 61 kPa to 263 kPa, between 20 mmHg and 150 mmHg for their models. Viscoelastic properties were neglected in this study due to the quasi-static nature of the inflation testing performed. Future work will focus on developing more realistic geometries by mimicking the dimensions of human carotid arteries.

Overall, this study presented a combined mechanical, US imaging, and computational framework. Fabricated homogeneous and bi-layered stenosed phantoms with lipid pool inclusions were explored using this framework improving our understanding of the biomechanical behavior of diseased vessels.

## 5. Conclusion

The study presents a framework combining inflation testing, US imaging and computational modelling of PVA phantoms. Bi-layered PVA vessels with different stenosis stages and lipid pool-like inclusions were successfully designed and pressurised. The results indicate the promising potential of PVA as a diseased artery surrogate to yield insights into the mechanical behaviour of vessels with atherosclerosis. Such experimental and computational work combined with non-invasive ultrasound imaging can provide a means to better understand why plaques are more prone to rupture and ultimately help improve the clinical management of carotid atherosclerosis.

## Acknowledgements

The authors would like to thank SIEMENS Healthineers for the ultrasound machine. This work was conducted with the financial support of the Science Foundation Ireland Centre for Research Training in Digitally Enhanced Reality (d-real) under Grant No. 18/CRT/6224. For the purpose of Open Access, the author has applied a CC BY public copyright licence to any Author Accepted Manuscript version arising from this submission.

## Notes

### Competing Interest Statement

The authors have declared no competing interest.

